# Mapping Molecular Gene Signatures Among Respiratory Viruses Based on Large-Scale and Genome-wide Transcriptomics Analysis

**DOI:** 10.1101/2021.10.17.464720

**Authors:** Thomas Smith, Mohammed A. Rohaim, Muhammad Munir

## Abstract

Severe acute respiratory syndrome coronavirus 2 (SARS-CoV-2) is an emerging RNA virus causing COVID-19 disease across the globe. SARS-CoV-2 infected patients exhibit acute respiratory distress syndrome which can be compounded by endemic respiratory viruses and thus highlighting the need to understand the genetic bases of clinical outcome under multiple respiratory infections. In this study, 42 individual datasets and a multi-parametric based selected list of over 12,000 genes against five medically important respiratory viruses (SARS-CoV-2, SARS-CoV-1, influenza A, respiratory syncytial virus (RSV) and rhinovirus were collected and analysed in an attempt to understand differentially regulated gene patterns and to cast genetic markers of individual and multiple co-infections. While a certain cohort of virus-specific genes were regulated (negatively and positively), notably results revealed a greatest correlation among gene regulation by SARS-CoV-2 and RSV. Furthermore, out of analysed genes, the MAP2K5 and NFKBIL1 were specifically and highly upregulated in SARS-CoV-2 infection in vivo or in vitro. In contrast, several genes including GPBAR1 and SC5DL were specifically downregulated in SARS-CoV-2 datasets. Additionally, we catalogued a set of genes that were conserved or differentially regulated across all the respiratory viruses. These finding provide foundational and genome-wide data to gauge the markers of respiratory viral infections individually and under co-infection.

## 1. Background

Since its first appearance in Wuhan, severe acute respiratory syndrome coronavirus 2 (SARS-CoV-2) has rapidly spread across the world in a way unlike any other respiratory viruses. Coronavirus disease 2019 (COVID-19) caused by SARS-CoV-2 is considered the third highly pathogenic coronavirus following SARS-CoV-1 and Middle East respiratory syndrome coronavirus (MERS-CoV) that cause severe accurate respiratory symptoms in humans [1]. The most striking feature of the incidences and epidemiology of SARS-CoV-2 is its high ability for transmission among people [2]. The clinical outcome and incidence vary that most COVID-19 patients are clinically mild and moderate, and the elderly seem to have serious symptoms [3]. Additionally, severely affected patients had shown respiratory complications such as moderate to severe pneumonia, acute respiratory distress syndrome (ARDS), sepsis, acute lung injury (ALI), and multiple organ dysfunction (MOD) [4].

ARDS in COVID-19 patients is thought to be the main cause of death because of the cytokine storm caused by an over-activation of the human innate immune response [5]. However, there are multiple immune regulators and host genetic and epigenetic factors that are capable of significant contributions to the disease manifestation [5]. Host-pathogen interactions can act as a double-edged sword in different coronavirus infections as they might be useful not just for hosts, but also for viruses [6]. Similar tug-of-war host-viruses can also be present in COVID-19, which could lead to overcomplicated outcomes of the disease [7].

Although recent studies have shown the transcriptomic analysis of host responses to SARS-CoV-2 infection at different time points within multiple cell lines [8, 9], the transcriptional dynamics of host response to multiple virus infection remained largely unknown. In general, the host innate immune responses play an essential role in suppressing the replication of the virus once the virus enters the host, such as antiviral-mediated interferons and cytokines, which could lead to the virus pathogenesis. Increased cytokine levels are also observed in patients hospitalised with COVID-19 in the same way as both SARS-CoV and MERS-CoV, which induce high levels of cytokine [10, 11]. Accordingly, understanding the magnitude and dynamics of human transcriptome in response to medically important respiratory viruses will help in understanding their pathogenesis, molecular genetic markers and in repurposing existing antivirals to combat respiratory viral infections.

The current study aims to compare a large cohort of transcriptomic dataset map the gene regulation (up or down regulated) by SARS-CoV-2 infection and the compounding impact of other respiratory viruses such as influenza, SARS-CoV-1, respiratory syncytial virus (RSV) and rhinovirus. This parallel comparison showcases common and unique genetic signatures of respiratory viruses under individual and co-infection scenarios.

## 2. Materials and Methods

### 2.1 Data Collection, Inclusion and Exclusion Criteria

Gene Expression Omnibus (GEO) and PubMed datasets were used to search for literature that contained data relating to upregulated and downregulated genes in response to infection with respiratory viruses (SARS-CoV-2, influenza, SARS-CoV-1, RSV and rhinovirus). The collection began with searching for datasets for the more recent COVID-19 pandemic. On GEO, the terms *“(“Severe acute respiratory syndrome coronavirus 2“[Organism] OR SARS-CoV-2[All Fields]) AND “Homo sapiens“”* were used whereas when searching on PubMed, the terms “*(SARS-CoV-AND (Transcriptome)”* were used. Once datasets were identified, inclusion and exclusion criteria were carried out as outlined in **Table 1** to ensure parallel comparison of gene signatures.

### 2.2. Included Datasets and Data Synchronisation

The collected datasets from various sources were compiled into one set of data using Microsoft Excel program. The studied viruses and their respective analysed datasets are provided in a spreadsheet (**Table 2**). An overview of each dataset is provided in the **Supplementary dataset 1**. Each dataset carried genes found in a specific study mentioned in the category, and the corresponding level of gene expression is displayed next units originally used by the datasets. To ensure that all the included datasets for each virus could be compared, these were converted to the same units. The raw data was often listed in three units; Fold Change, Log Fold Change and Log 2-Fold Change, and all the data was converted into the Log 2-Fold Change format. Log 2-Fold Change was used as it allows easier visualisation of the data, as the range of the values of the data becomes narrower, allowing for easier comparison of the up/ down regulated genes **(Supplementary dataset 2)**.

### 2.3. Ranking System

Owing to large diversity among datasets in areas such as cell types and media in which the experiments were carried out, it could introduce biasness to compare genes specifically by their Log 2-Fold Change values, which is calculated to the baseline gene expression. To introduce a novel method of comparing each gene up or down regulated in a dataset compared to datasets from another viruse or different cell types, a ranking system next to each Log 2-Fold Change column was proposed. This system ranked the genes based on which percentage group they were in, depending on whether they were up or down-regulated. Then, a mean score was taken across datasets within the same studied viruses and these means were used to compare between the viruses. For avoidance of confusion, this system synchronizes the dataset such that at the top 10% of upregulated genes for one virus while only at the top 80% of genes for another virus.

Using the GraphPad Prism 9.0.0 software, a scatter bar graph was generated using the overall ranking score for each gene of each virus. Two versions were created; first had the uncut data taken directly from the ranking system, containing roughly 24,000 genes, and secondly a cut down version of the data where non-significant genes were removed. Additionally, the non-coding gene loci and non-annotated genes were removed, as these often yielded zero values for up or down regulated genes reducing 6200 genes. Furthermore, other genes were removed which contained more than three or more zero values for up or downregulation across the five viruses removing a further 200 genes. Finally, using influenza virus as a model virus, all genes were removed that lied within the ranking scores of +20 (bottom 20% upregulated) and −20 (bottom 20% downregulated genes), unless a gene had a ranking score of above +50 or below −50 in any other of the viruses. This removed a further 5005 genes leaving a total of just over 12,000 genes in the cut down version, which removed the large proportion of genes containing zero values for clearer view for the spread of gene ranking scores.

### 2.4. Log2 Fold Change

The collected dataset was converted into Log2 Fold Change for the gene expression. The datasets that were unconvertable into Log2 were removed for Log2 Fold Change analysis. In addition, only datasets that compared infected and non-infected patients were used while high vs low viral load datasets were removed. Finally, Log2 Fold Change values for each gene and for each dataset were inputted into the software and the graphing tool was used to generate scatter bar charts for each virus-specific dataset. These include the top five upregulated and/or downregulated genes for each dataset taken from the original data.

### 2.5. iDEP.91 Software

Once all the data had been converted into the ranking score format, it was exported into a separate Excel File to be compiled into one concise table **(Supplementary dataset 3)**, then saved as a text document and uploaded to the iDEP web application for expression and pathway analysis as described earlier [12].

## 3. Results

### 3.1. Overview of the differences in the Log-2 Fold Change values and Ranking Scores Across Multiple Respiratory Viruses

The scatter bar graphs for each of the individual datasets within each of the five viruses were drawn to provide an overview of the differences in the Log-2 Fold Change values obtained from each dataset (**Fig. 1)**. The scatter bar graph for the datasets collected for the SARS-CoV-2 uses the original Fold change values given by each study where each bar represents a separate dataset that showed the up and down regulated genes in response to viral infection (**Fig. 1A)**. A vast majority of top five upregulated genes were summarized (**Table 3**) while the top five down regulated genes involved in the innate immune response to SARS-CoV-2, SARS-CoV-1, influenza, RSV and rhinovirus infection were concluded (**Table 4**). Interestingly, each dataset shown was distinctive showing a varying pattern where host genes are mildly up or down regulated and only a few that are highly differentially up or down regulated. This highlights selective genes of the innate immune response are affected in response to a specific virus infection. Collectively, dataset GSE155286 has the widest spread of data while dataset GSE147507 has the lowest **(Table 2)**. In addition, all the datasets arried both downregulated and upregulated genes except GSE153790, which has only upregulated genes. Amongst the top five up-regulated genes, five genes including IFI27 and C-X-C motif chemokine ligand (CXCL) group of cytokine-producing genes, specifically CXCL10 showed a virus-specific trend.

**Fig. 1.**
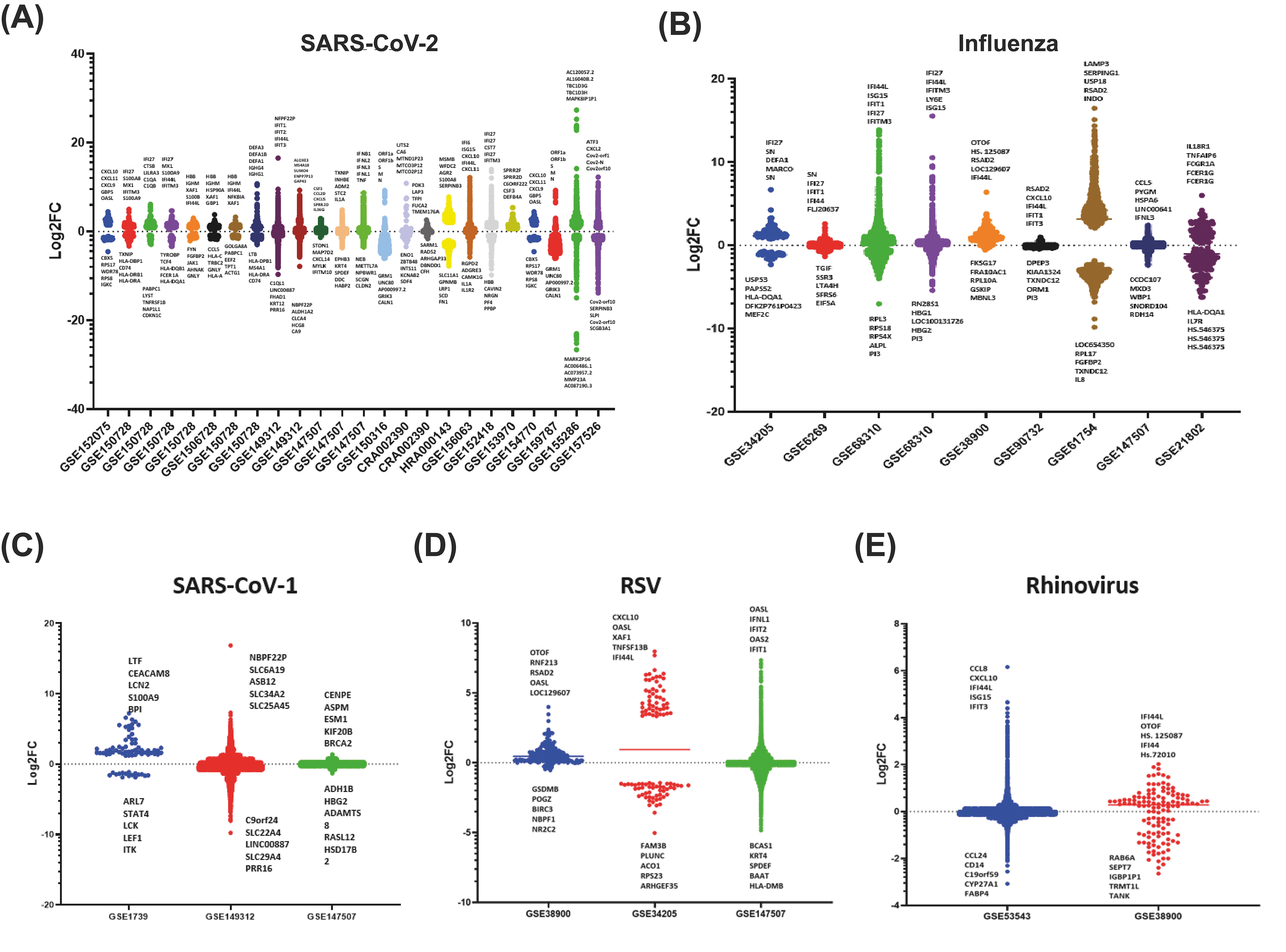
Scatter bar graphs of the Log-2-Fold Change of each gene for each dataset for the A) SARS-CoV-2, B) Influenza, C) SARS-CoV-1, D) RSV, E) Rhinovirus. A horizontal line is also shown on each bar, which marks the average Log-2 fold change of the selected genes.

Using the same approach, we used the data collected for innate immune genes in response to influenza virus infection that contains nine datasets. The data was presented for better visualising to gauge the innate immune genes play critical roles in the virus infection. Consistently, amongst all datasets, the up regulated genes for the influenza virus were interferon alpha-inducible protein 27 (IFI27) and interferon induced protein 44 producing gene IFI44/IFI44L, which involves in type-1 interferon signalling process leading to apoptosis and the formation of tubular structures, respectively.

The scatter bar graphs for SARS-CoV-1, RSV and rhinovirus indicate a unique set of genes up or down regulated during infection (**Fig. 1C, 1D and 1E)**, respectively. While limited datasets were available against some viruses, minimum eight datasets provided approximately 12,000 different genes. Datasets that have gaps around the zero value for Log-2 fold change are the datasets that only include genes that were significantly up or down regulated. All datasets shown in **Fig. 1C, 1D and 1E** show a clear abundance of genes that are mildly differentially regulated with significantly less genes at the high fold change values, highlighted by the shape of the GSE53543 dataset. Interestingly, there was marked variation between the highest and lowest values obtained for log-2-fold change for different datasets within SARS-CoV-1. In addition, most of innate immune genes fall within +10 or −10 log-2 fold change for these viruses. However, SARS-CoV-1 appears to have a unique set of top five up regulated genes compared to the other viruses whereas both RSV and rhinovirus datasets showed IFI44 gene and the CXCL family. OASL remained a consistently upregulated gene in RSV datasets.

The log-2 fold change values of each gene for each dataset was changed into a ranking score due to the high variation of experimental method used to collect data for each dataset, which meant that log-2 fold change values were rarely consistent between datasets for differential gene regulation of patients/cells infected with the same respiratory virus. Thus, the ranking score removed this issue by assigning each gene a value based on its position among other differentially regulated genes within the same dataset (i.e., a gene placing as the 5^th^ highest upregulated gene in a list of 100 genes would receive a score of 95). These synchronized values were averaged across all datasets within each virus that enabled the data collected from different experimental approaches to be compared more effectively between datasets within the same virus and a combination of datasets to be compared between different viruses **(Table 5 and 6)**.

### 3.2. iDEP.91 statistical analysis

The application of ranking scores facilitated the generation of a dataset consisting of 12,000 genes across all viruses by removing many non-significant genes (**Fig. 2A**). This newly and reduced set of genes and the data provided a higher resolution of genes distribution across multiple respiratory viruses (**Fig. 2B)**. Thereafter, all analysis was conducted using dataset generated through ranking system

**Fig. 2.**
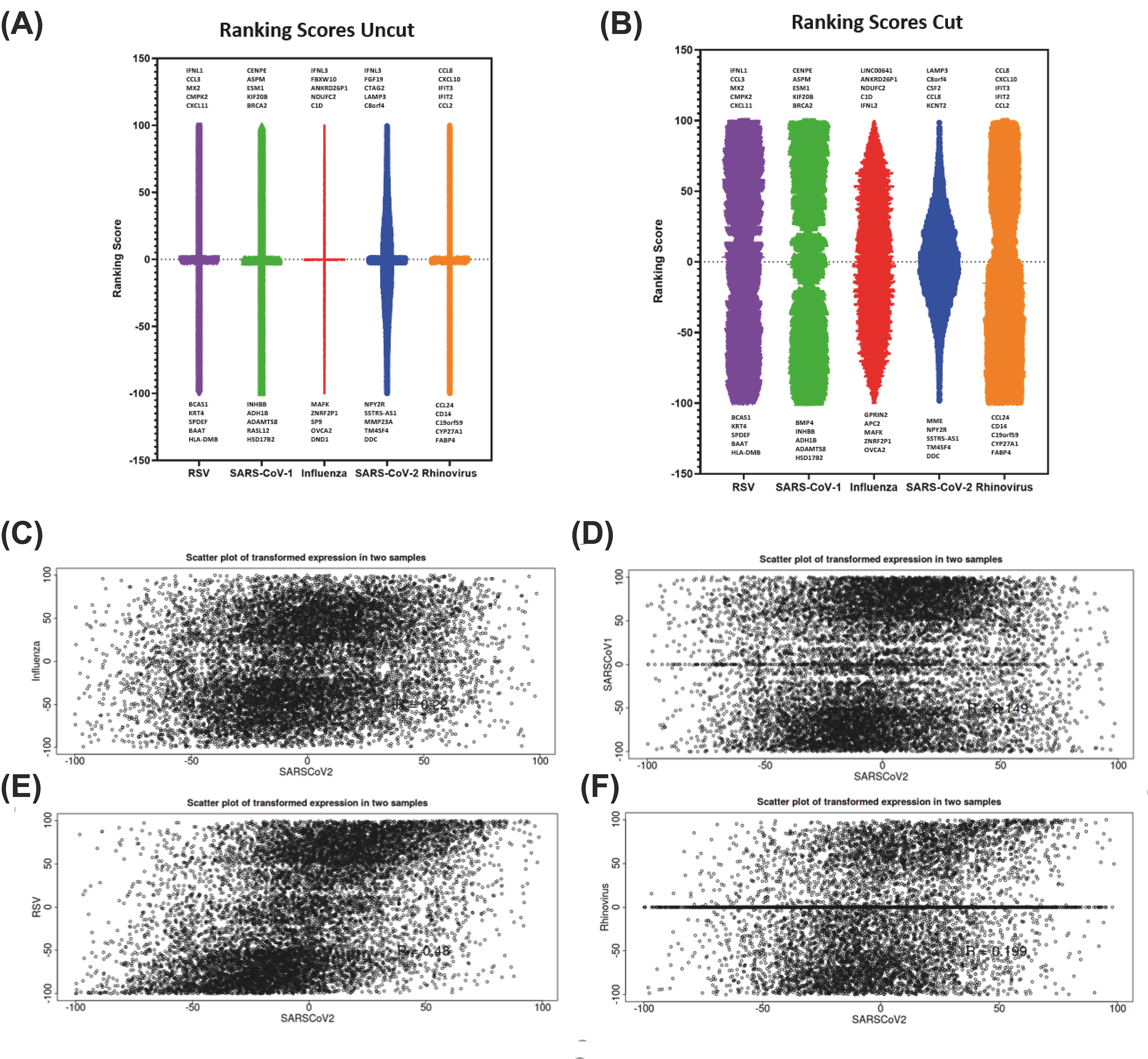
Uncut (A) and Cut (B) ranking scores for each gene combining all datasets for each respiratory virus. Also, in this figure are scatter plots of ranking scores of all genes collected for each respiratory virus, using SARS-CoV-2 as the comparison. (C) shows a comparison of Influenza and SARS-CoV-2, (D) between SARS-CoV-1 and SARS-CoV-2, (E) between RSV and SARS-CoV-2 and (F) between Rhinovirus and SARS-CoV-2.

The scatter plots generated on 12,000 genes highlight the distribution patterns of genes between SARS-CoV-2 and other respiratory viruses (**Fig. 2C to 2F)**. The relationship between SARS-CoV-2 and influenza virus gene regulation revealed a uniform scatter data (**Fig. 2C)**, while the relationship between SARS-CoV-2 and SARS-CoV-1 gene regulation contains more spread of data points except towards the centre of the graph due to the removal of less important data towards zero values **(Fig. 2D**). A slightly different patterns was observed where a linear relationship between SARS-CoV-2 and RSV (**Fig. 2E**) was noticed. An overall less uniform spread of data points with a skew to the right towards the top of the graph, and additional upregulated genes were observed in SARS-CoV-2 and rhinovirus comparison (**Fig. 2F)**.

### 3.3. Heatmap Analyses and Gene Differences between Respiratory Viruses

The heatmap were generated to provide an insight into pathways that are differently regulated by each of the five studied respiratory viruses (**Fig. 3)**. SARS-CoV-2 appeared unique in eliciting a separate viral response compared to the other respiratory viruses. Notably, there was a region at the bottom of the heatmap between genes DDX21 and GBP3 where other viruses have no effect or a slight upregulation of the genes, however, SARS-CoV-2 causes a downregulation (**Fig. 3)**.

**Fig. 3.**
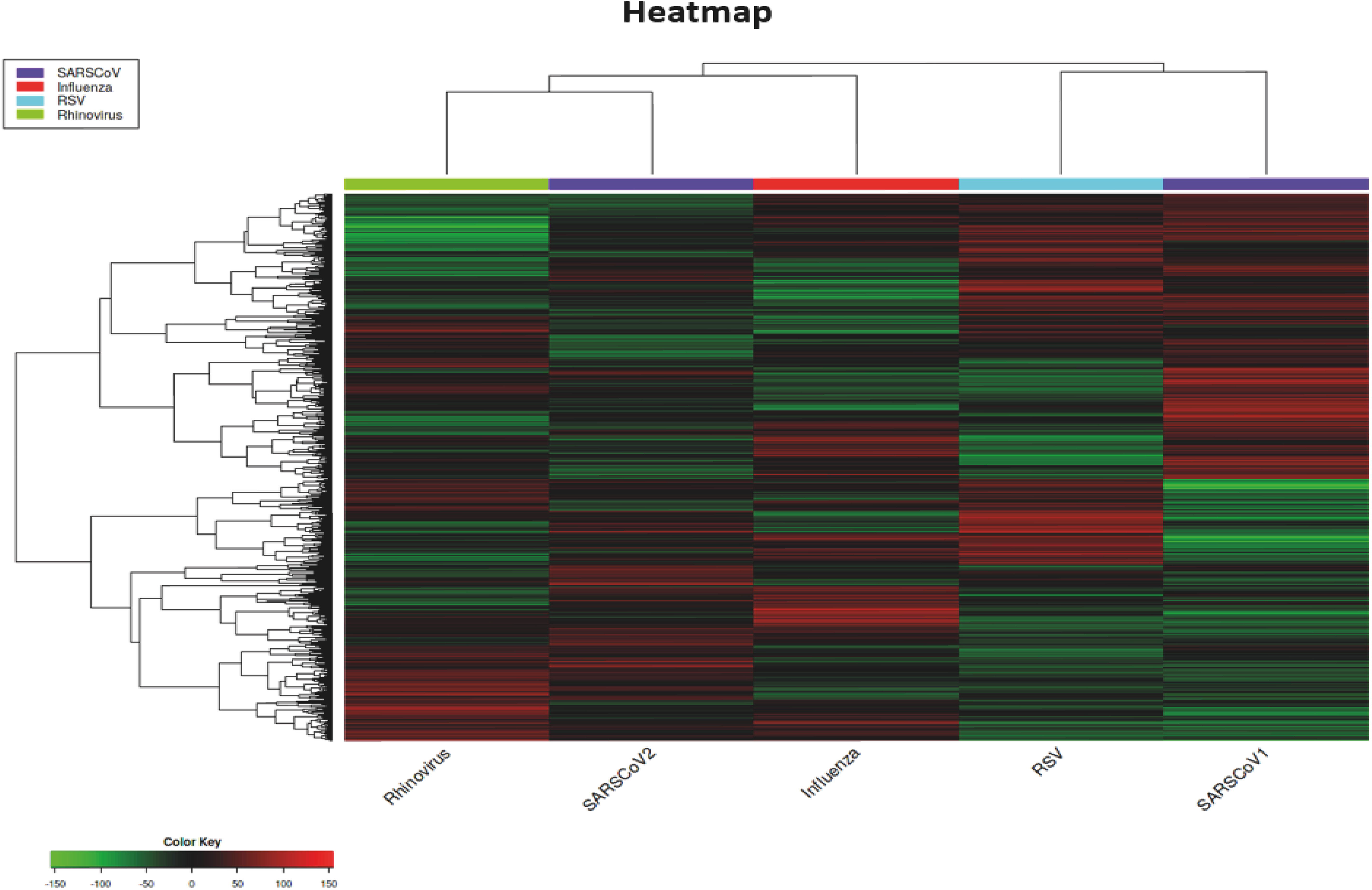
Heatmap of DEGs for all the respiratory viruses studied in this analysis.

Perhaps the most unique out of all the respiratory viruses is SARS-CoV-1 which showed large areas of each heatmap where it is causing a downregulation of genes where all other respiratory viruses were eliciting upregulation.

The heatmap results highlighted the differences between each of the respiratory viruses, even though they are in the same group based on their target within the host; the genes that being affected are substantially different. Each virus shown in the heatmap carried different and distinct green and red areas, with very few coloured areas shared between more than two viruses. The most substantial difference was noticed between SARS-CoV-2 and SARS-CoV-1, whereas almost no colours in common. However, SARS-CoV-1 appeared to be the only virus that has both up and down regulated genes in two specific groups.

### 3.4. Standard deviation calculation and T-SNE plot Analyses

The SD graph highlights the extremely high standard deviation across all the regulated genes in response to different viruses (**Fig. 4A)**. A standard deviation above 1 was considered high unless the standard deviation in this case was between 25 and 75 indicating that there are high differences in the differentially regulated genes in response to each virus. On the other hand, a correlation matrix that shows the correlation between each of the viruses revealed that the most similar virus to SARS-CoV-2 was RSV with a Pearson’s correlation coefficient of 0.48 **(Fig. 4B)** while the least similar one was SARS-CoV-1 with a Pearson’s correlation coefficient of 0.15 **(Fig. 4B)**. A correlation value of 1 implies that there is a perfectly linear distribution of data between the two variables and a value of 0.48 generated for RSV compared to SARS-CoV-2 is relatively high that highlight how close the two viruses are in comparison to other viruses.

**Figure 4.**
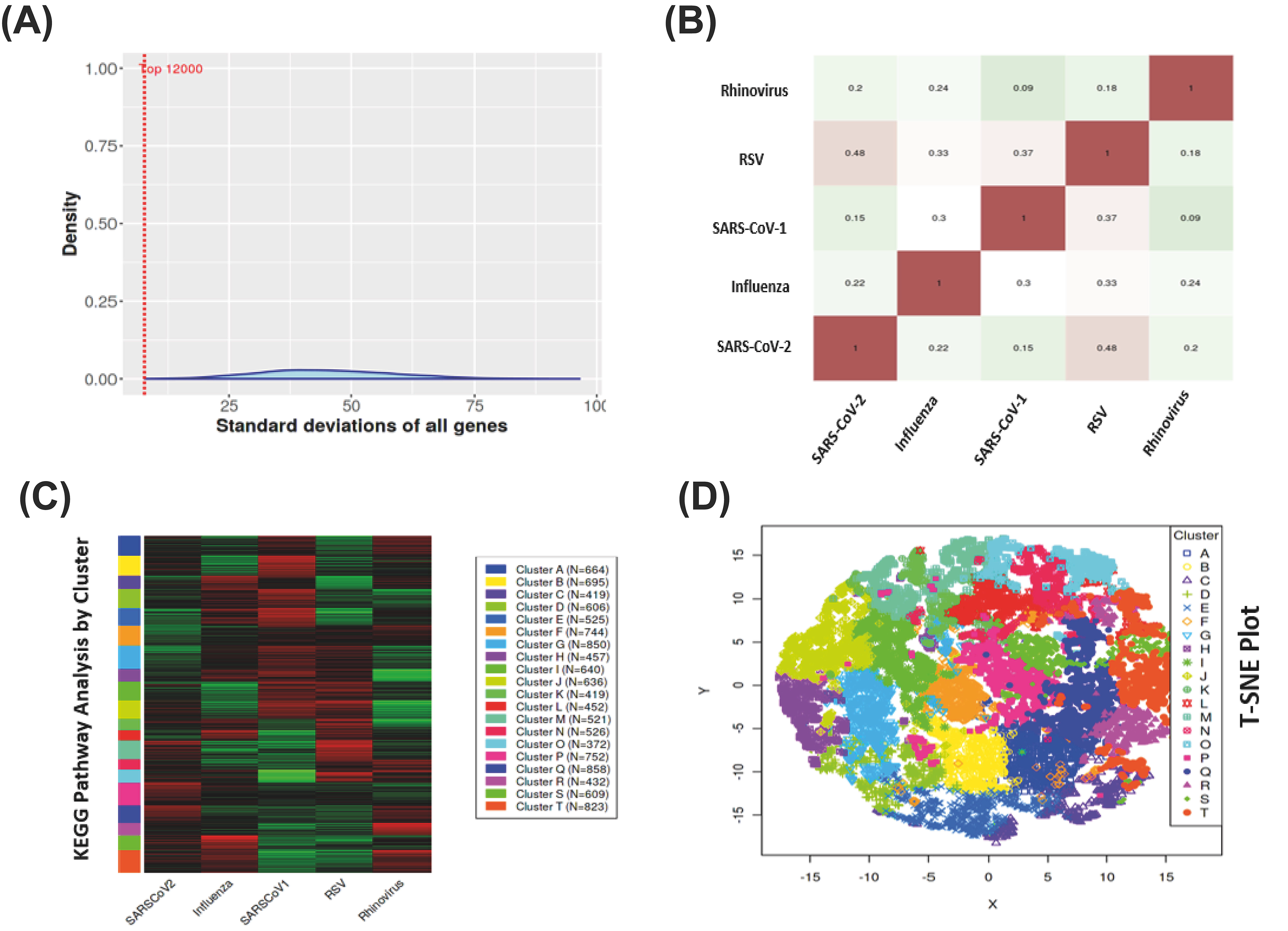
(A) Standard deviation of all genes across all viruses. (B) Correlation matrix using data taken from the top 75% of genes. (C) KEGG pathway analysis by cluster. (D) T-SNE plot of all 12,000 genes

Differentially regulated genes were classified into 20 clusters based on their K means (**Fig. 4C)** where we used them to break down for better understanding whereabouts the differences between these emerged viruses. Each cluster contains genes involved in specific pathways that allows for the comparison of gene regulation in a variety of pathways depending on the virus **(Supplementary dataset 4)**. After K-means clustering, cluster O appeared to contain the most pathways involved in the innate immune response, such as the JAK-STAT signalling pathway, TNF signalling pathway and IL-17 signalling pathway indicating that cluster O could be used as a sign of a virus’s regulation for the overall innate immune response signalling. Both influenza and SARS-CoV-2 showed both up and down regulated genes within the cluster with specific areas either being highly up- or down-regulated, suggesting that these viruses target specific areas within this cluster. While SARS-CoV-1 and RSV upregulated and down regulated this region, respectively.

The T-SNE plot analyses for all the data was coloured based on their belonging cluster. The T-SNE allowed multi-dimensional data to visualise in a low dimension space such as the 2D graph **(Fig. 4D)**. The distance between each of the points reflected the similarity of each data point. Whilst T-SNE should not always be used for gene expression data analysis, due to its high intrinsic dimensionality. Therefore, it has been used to highlight that even though there are a high number of clusters present, they are still very much distinguishable, despite there being some clusters that exhibit more separation of data points compared to others. In addition, there was a slight problem with crowding towards the centre of the dataset; however, this was observed in most SNE forms.

### 3.5. Comparison between differentially regulated genes among multiple respiratory viruses

Generally, the number of upregulated genes is relatively even with the number of downregulated genes, however, there are more downregulated genes than upregulated genes for each of the five tested viruses. The standouts are substantially downregulated than upregulated genes in case of rhinovirus infection in (**Fig. 5A)**. Moreover, rhinovirus showed less differentially regulated genes in total compared to the other respiratory viruses.

**Fig. 5.**
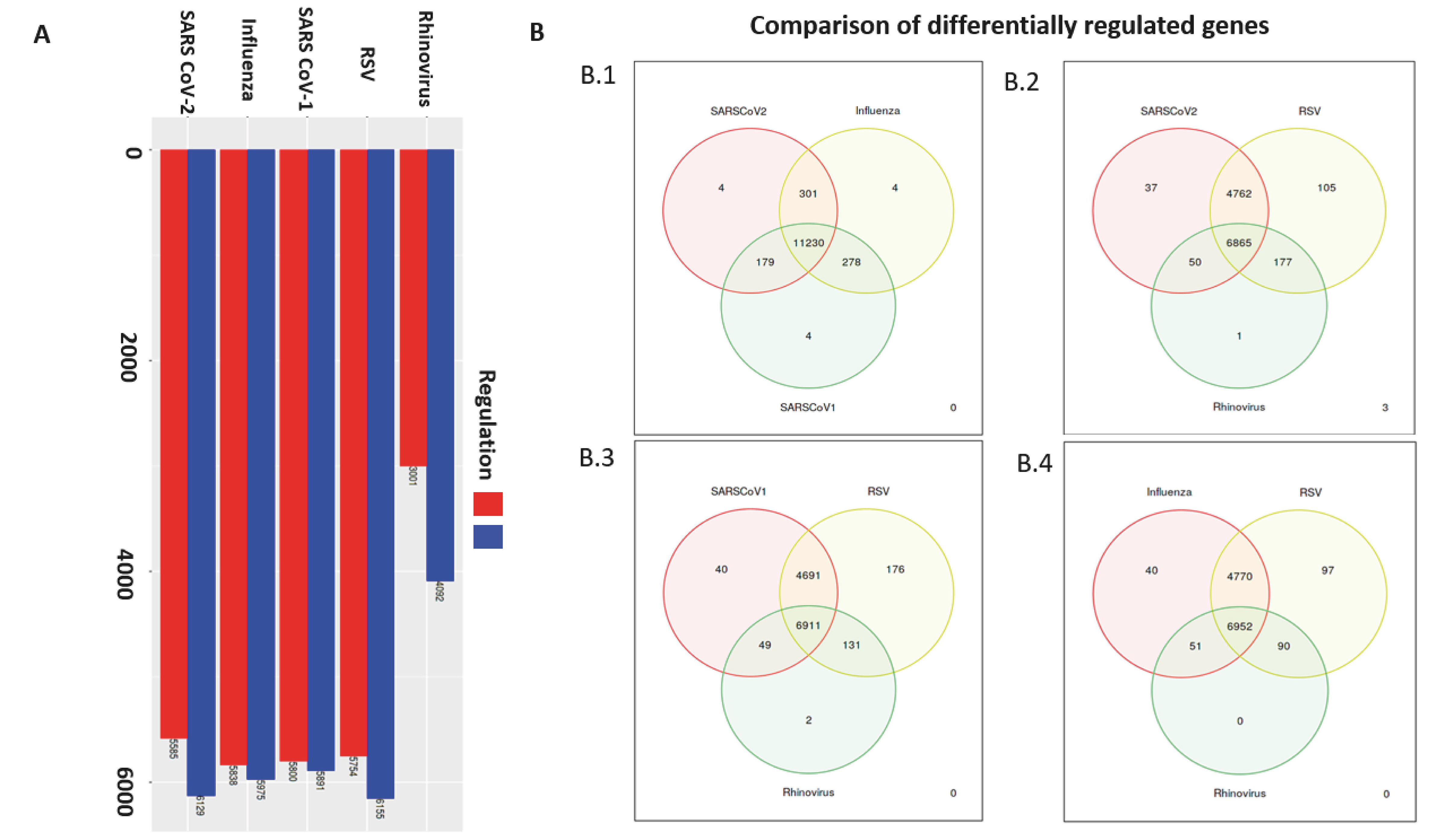
(A) Total number of upregulated and down regulated genes for each virus. (B) Venn diagrams representing the differentially regulated genes that are in common between each of the respiratory viruses.

The Venn diagrams showed a comparison between each of the viruses by how many differentially regulated genes they have in common, regardless of whether they are up or down regulated. This highlighted genes that are differently regulated within only one virus compared to others within the same diagram (**Fig. 5B**). Vast majority of genes are found to be differentially expressed across all viruses; however, there were some exceptions mainly found within RSV that has the highest number of genes unique to itself while rhinovirus rarely had any uniquely expressed genes.

### 3.6. Impact of SARS-CoV-2 on cellular DNA replication

A visual representation for the impact of SARS-CoV-2 on the DNA replication within infected cells was outlined (**Fig. 6)**. The upregulated genes (2/32) were shown in red while the downregulated (27/32) were shown in green **(Fig. 6)**. Genes responsible to produce DNA ligase and helicase were notably down regulated, which are important in the DNA replication and being used by the virus as a means of slowing down the cell cycle to enhance viral replication.

**Fig. 6.**
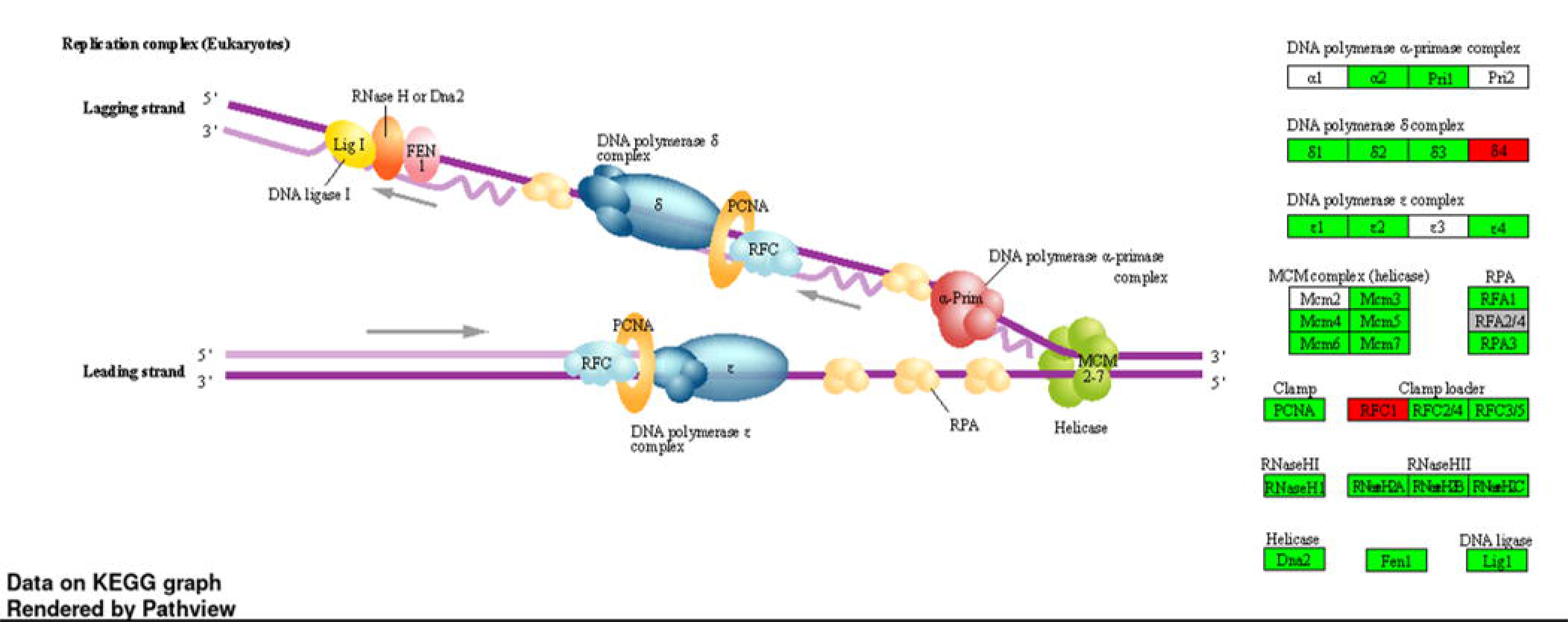
Heatmaps specific to different pathways compiled by GAGE pathway analysis. (A) for Defence response to virus, (B) for cytokine response, (C) for regulation of cytokine production and (D) for positive regulation of innate immune response.

### 3.7. Regulation of JAK-STAT immune signalling pathway in response to SARS CoV-2 infection

There are more upregulated genes in JAK-STAT immune signalling (**Fig. 7A)** and the cytokine-cytokine receptor interaction pathways (**Fig. 7B)** than downregulation, highlighted by the prominence of the red colouring over the green colouring. While there were several upregulated genes as GFAP and Ras, which are involved in cell differentiation and MAPK signalling pathway, respectively.

**Fig. 7.**
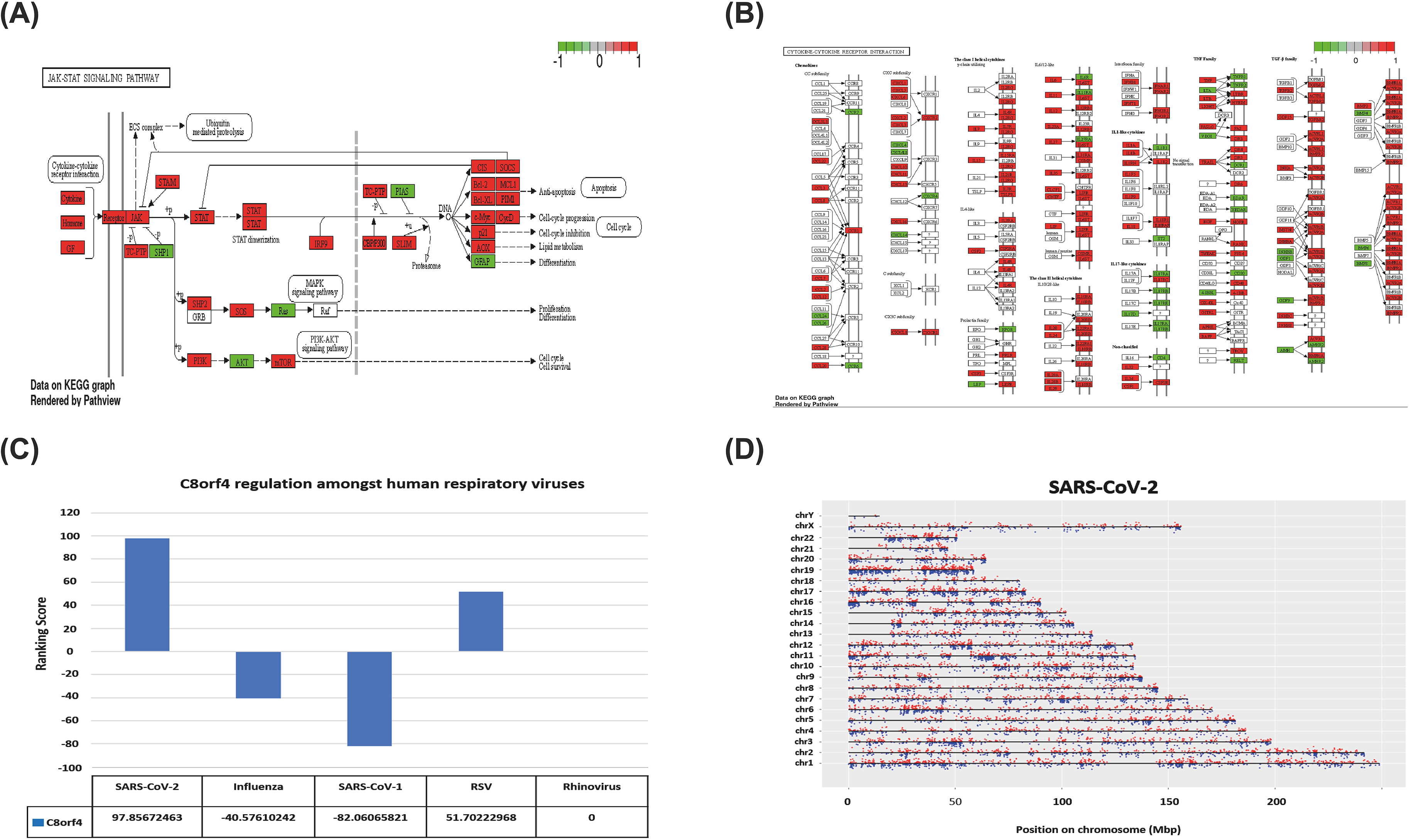
(A) Regulation of genes associated with the JAK-STAT signalling pathway. (B) Regulation of genes associated with cytokine-cytokine receptor interaction. (C) Ranking scores of the C8orf4 gene for each respiratory virus. (D) Genome map showing SARS-CoV-2 upregulated genes in red and downregulated genes in blue.

The genes that are up or down regulated in realtion to the immune signalling pathways and are affected in response to SARS-CoV-2 infection were analysed using KEGG pathway database (**Fig. 7A, 7B and Supplematary Figure 1)**. These results revealed that SARS-CoV-2 does not affect every pathway in a simple manner by either upregulating or downregulating all genes involved in that pathway, but instead having multiple effects.

Using the ranking scores, C8orf4 was the second most highly upregulated gene in cells/patients infected with SARS-CoV-2. The C8orf4 (also known as TCIM) is responsible for producing the c8orf4 protein (also known as TC1) which is involved in the enhancement of NF-kappaB activity, leading to up-regulating several cytokines involved in the process of inflammation [13]. This is the main factor attributed to the cytokine storm exhibited in patients following SARS-CoV-2 infection. In addition, our analyses show that each virus has a different effect on the regulation of c8orf4 and its regulation could therefore be used as a biomarker to differentiate between aetiology of infection, with extremely high levels of TC1 protein pointing towards a SARS-CoV-2 infection **(Fig. 7C)**. Of course, many other genes could be used as markers for SARS-CoV-2 infection but also genes that are conserved between all viruses.

After individual identification of the upregulated or downregulated genes and their respective pathways, we aim to visualise where those genes are located within the human chromosome **(Fig. 7D)**. Human genome map analyses show each chromosome with its own line with genes where the upregulated genes appear above the line in red colour while genes that are downregulated appear below the line in blue colour **(Fig. 7D)**. This genomic map shows the regulation in response to SARS-CoV-2 infection and revealed that every chromosome in the human genome has been affected whereas the mostly affected chromosome was chromosome 19. However, the least affected chromosomes were X and Y sex chromosomes. In addition, chromosome 17 also shows a notable pattern. There are many areas across many chromosomes that showed notable gaps where SARS-CoV-2 appears to have no effect on gene regulation **(Fig. 7D)**.

There is a large amount of consistency between all the genome maps within the most affected chromosome, in all cases, being chromosome 19. In case of rhinovirus, there is a lack of altered genes regulation on the X and Y chromosomes. Furthermore, a much blander overall picture on fewer data points (**Supplementary figure 2D**) because there were less genes recorded to have been up or downregulated in the rhinovirus dataset.

## 4. Discussion

Despite majority of the human respiratory viruses show similar pathology by infecting the same respiratory system, they all showed clear and substantial differences, which have highlighted unique markers related to differential gene regulation. The scatter plots showed the correlation between the effects of each virus on human gene expression, and a specific removal of genes was evident in this analysis which are less dramatically differentially regulated and therefore of less importance to this study. These results indicated that SARS-CoV-2 is like RSV compared to other respiratory viruses because of the high correlation between the data points within the scatter graphs showing a rising diagonal line suggesting a positive correlation between the upregulated and downregulated genes. These results are supported by the correlation matrix, where the Pearson’s correlation coefficient between SARS-CoV-2 and RSV was 0.48, much higher than the 0.22, 0.2 and 0.15 for influenza, rhinovirus, and SARS-CoV-1, respectively. The SARS-CoV-2 and RSV showed high similarity in differentially regulated genes. This aligns with the fact that affected patients exhibit similar symptoms when infected with any of SARS-CoV-2 or RSV, mainly upper respiratory tract symptoms and often lower respiratory tract symptoms such as a dry cough [14]. Interestingly, both viruses appear to cause damage to the respiratory tract that result in persistent problems long after infection such as persistent airway obstruction as well as hyper-responsiveness can be seen in patients 30 years after infection with RSV [15]. These symptoms are like the long-term lung dysfunction reported after SARS-CoV-2 infection [16]. However, the main difference between these two viruses is the age of the patients that are more susceptible for infection, with RSV commonly causing respiratory tract infection in young infants and children [14], whereas SARS-CoV-2 is known for more severe cases being present in the elderly albeit infection potential among all ages. Further research in this area could be useful to compare influenza, RSV, SARS-CoV-1 and rhinovirus against SARS-CoV-2 but specifically for each pathway/area such as the innate immune response or the cytokine activation pathway.

Insights into the human chromosomes in response to SARS-CoV-2 infection revealed that the mostly affected chromosome was chromosome 19 suggesting a high number of genes involved in the immune response to viral infection could be present within chromosome 19 and severe cases of infection could be attributed to the genetic mutations within this chromosome. Another interesting point is the presence of differential gene expression on the X chromosome for patients suffering from COVID-19. Altered genes on the X-chromosome could lead to a difference in the clinical outcome between men and women infected with SARS-CoV-2. Previous studies reported that the immune regulatory genes encoded by the X chromosome in women could cause lower viral load levels resulting in a reduction in the inflammatory response compared to men [17].

The top and bottom five consistently up and down-regulated genes across all five viruses could potentially be used as markers for specific respiratory viral infection. JAK2 is one of the genes, which is consistently, and highly upregulated among all the studied viruses and it encodes for the Janus Kinase 2 protein (JAK2). JAK2 plays a crucial role in the cytokine signalling where it associates with type II cytokine receptors, hormone-like cytokine receptors and being activated by IFN-gamma [18]. Additional four-upregulated genes were DDX60L, IFI44, FOXN2 and DDX60, which may be a target for drugs.

The upregulated genes in response to SARS-CoV-2 infection have been identified while those were downregulated in the other respiratory viruses. These genes could be used as markers for a SARS-CoV-2 infection and to distinguish SARS-CoV-2 from other respiratory viruses. The most important gene was the NFKBIL1 gene that encodes for the NF-kappa-B inhibitor-like protein 1 and it is involved in the NF-kappa-B signalling, which plays a major role in the inflammatory response by increasing the cytokine expression [19]. On the other hand, the downregulated genes in response to SARS-CoV-2 could possibly be used as a marker to distinguish SARS-CoV-2 infection in case of suspicion with a respiratory virus infection associated with respiratory symptoms. One of these genes is GPBAR1, which encodes for the G-protein acid receptor 1. Previously, it has been reported that GPBAR1 was able to regulate and increase the expression of IL-10 [20] suggesting that levels of IL-10 in patients suffering with COVID-19 would be lower, however, recent studies contradict that as IL-10 levels are found to be unexpectedly increased in severe cases [21].

## 5. Conclusions and limitations

The aim of this study was to determine the influence of SARS-COV-2 on the immune regulation and gene induction in comparison to other respiratory viruses. It appeared that SARS-CoV-2 was unique in its impact on gene regulation and matches none of the other respiratory viruses except RSV. Genes such as MAP2K5 and NFKBIL1 have been found to be greatly upregulated in SARS-CoV-2 whilst being downregulated in the compared viruses. Whereas genes such as GPBAR1 and SC5DL were contrastingly found to be significantly downregulated in SARS-CoV-2 but upregulated in influenza, SARS-CoV-1, RSV and rhinovirus. Despite all the reported differences, the most conserved genetic signature was JAK2 gene as well as the constitutively downregulated ZNF219 gene. While the resolution of analysis provides foundational finding, further research is warranted to validate the impact of these molecular signature against individual or multiple infections.

We observed a limitation of the study that the gene regulation may be affected by the experimental characteristics such as time length post infection, the culturing conditions, phenotypes of the cells, and the nature of the virus stimulation (in vivo or in vitro studies). Finally, different cell types (A549, BALF or PBMC cells) were used for virus infection, which may respond differently to different viral infections. Nevertheless, the provided analysis provides a foundation for the impact of respiratory viruses on the gene regulation.

## Ethics Statement

No experimental procedures were carried out in this project and all data was collected from a range of previous research papers; therefore, no steps of this study were required to seek ethical approval.

